# Metastasis-specific gene expression in autochthonous and allograft mouse mammary tumor models: stratification and identification of targetable signatures

**DOI:** 10.1101/2019.12.20.885210

**Authors:** Christina Ross, Karol Szczepanek, Maxwell lee, Howard Yang, Cody J. Peer, Jessica Kindrick, Priya Shankarappa, Zhi-Wei Lin, Jack Sanford, William D. Figg, Kent Hunter

**Affiliations:** Laboratory of Cancer Biology and Genetics, Metastasis Susceptibility Section, Center for Cancer Research, National Cancer Institute, Bethesda, MD, USA; Laboratory of Cancer Biology and Genetics, High-Dimension Data Analysis Group, Center for Cancer Research, National Cancer Institute, Bethesda, MD, USA; Clinical Pharmacology Program, Office of the Clinical Director, National Cancer Institute, NIH, Bethesda, Maryland

## Abstract

Breast cancer is a leading cause of cancer-related death of women in the U.S., which is ultimately due to metastasis rather than primary tumor burden. Therefore, increased understanding of metastasis to develop novel therapies is critical in reducing breast cancer-related mortality. Indeed, a major hurdle in advancing metastasis-targeted intervention is the genotypic and phenotypic heterogeneity that exists between primary and secondary lesions. To identify targetable metastasis-specific gene expression profiles we performed RNA sequencing of breast cancer metastasis mouse models. We analyzed metastases from models of various oncogenic drivers and routes, including orthotopic injection, tail vein injection, intracardiac injection, and genetically engineered mouse models (GEMMs). Herein, we analyzed samples from 176 mice and tissue culture samples, resulting in 338 samples total. Using these data, we contrasted the different breast cancer metastasis models, and also identified common, targetable metastasis specific gene expression signatures.

Principal component analysis revealed that mouse model (Allograft v. GEMM) rather than tissue source (PT v metastatic nodule) shaped the transcriptomes of our samples. Allograft models exhibited more mesenchymal-like gene expression than GEMM models, and primary culturing of GEMM-derived metastatic tissue induced more mesenchymal-like gene expression. Furthermore, metastasis-specific gene expression differed between tail vein and orthotopic injection models of the same cell line, the degree of which was cell line dependent. Finally, we examined metastasis-specific gene expression common to models of spontaneous metastasis (orthotopic injection and GEMMs). Pathway analysis identified upregulation of the sildenafil response, and nicotine degradation pathways. The influence of these pathways on metastatic spread was assessed by treatment of allograft models with clinically relevant dosing of sildenafil or nicotine. Sildenafil significantly reduced pulmonary metastasis while nicotine administration significantly increased metastasis, and neither regimen altered primary tumor mass. Taken together our data reveals critical differences between pre-clinical models of metastatic breast cancer. Additionally, this strategy has identified clinically targetable metastasis-specific pathways integral to metastatic spread.

## Introduction

Metastatic breast cancer remains the leading cause of cancer-related death among women world-wide^1^. Standard treatment for breast cancer patients diagnosed with stage I-III is surgical resection, indicating that mortality is largely due to recurrence at distant sites^2^. Indeed, while non-metastatic breast cancer has a 5-year survival rate of 98.8%, metastatic disease reduces 5-year survival to only 27.4%^1^. Therapeutic strategies to treat localized disease, such as molecular profiling and targeted therapy, have been increasingly successful, but patients with disseminated disease continue to face much worse outcomes, as metastases are largely insensitive to current treatments^3,4^. Therefore, to improve outcome for patients with advanced cancer, specific metastasis-targeted strategies will need to be developed, as will a deeper understanding of the unique biological processes that occur during disease progression^4–6^.

Investigations of metastatic biology trail those of primary tumors (PTs) due to difficulty in obtaining appropriate human tissue samples. Metastatic lesions are usually not surgically removed, and those that are resected are usually confounded by treatment^7^, which may enrich for characteristics associated with therapy resistance rather than metastasis. Additionally, early biopsies of metastatic lesions in untreated patients often have low tumor cell content^7,8^ or are not accessible to biopsy, and repeated biopsies of lesions within individuals increases patient morbidity^9^. As an added challenge, since metastases within patients exhibit significant heterogeneity at both the biomarker^10–12^ and genomic levels^13,14^, selective biopsies may not capture the full complexity of events associated with tumor progression. Therefore, although human data from smaller genomics projects focused on metastasis are now becoming available^15,16^, our understanding of the somatic events that contribute to metastasis remains far behind the understanding of primary tumorigenesis.

Mouse models overcome many of these limitations and offer additional advantages: (i) a readily available, renewable resource of tissue enables investigation of a genetically equivalent set of tissues over time; (ii) the ability to observe the full natural history of metastatic development, which is not possible in patients due to surgical resection and application of neo-adjuvant and adjuvant therapies; (iii) the existence of syngeneic cell line injection (allograft) models to rapidly test hypotheses generated from genomic characterization; and (iv) genetically engineered models to examine factors that function through somatic microenvironmental effects rather than directly within tumor cells, an aspect often precluded in human studies. Data generated from animal models therefore provide an important vehicle for hypothesis generation and testing that can subsequently be examined and verified in more restrictive human experimental systems.

As mouse models have become integral to the study of metastatic spread, the variety of pre-clinical mouse models so too has grown to better understand the steps of the metastatic cascade and capture the many stages of this disease. To identify metastasis-specific gene expression (MSGE) profiles that are integral to the establishment and growth of metastatic lesions, we performed RNA-sequencing (RNA-seq) of breast cancer metastases and paired primary tumors isolated from several mouse models. As the interventions for metastatic cancer are limited to targeting already established secondary tumors^17^, we analyzed primary tumors and macrometastases from models of various oncogenic drivers and routes, including orthotopic injection, tail vein injection, intracardiac injection, and genetically engineered mouse models (GEMMs). Here we present data revealing critical differences in MSGE between pre-clinical models of metastatic breast cancer, important differences that we anticipate will shape the use of these models by the field moving forward. Additionally, we have identified several core and targetable gene expression pathways common to multiple models, which we further confirmed the significance of *in vivo*. By combining the analysis of tissues from several models of metastasis we present here a robust strategy for the discovery of clinically targetable pathways that may be integral to metastatic spread.

## Results

### Significant transcriptional differences exist between mouse models of metastatic breast cancer

To investigate the transcriptomic landscape specific to metastatic breast cancer, RNA-seq analysis of the most commonly used mouse models of metastatic breast cancer was performed. Matched primary and metastatic tumor tissues from GEMMs of metastatic breast cancer were collected from four mouse models representing luminal (MMTV-Myc, MMTV-PyMT), basal (C3(1)-TAg), and HER2+ (MMTV-Her2) subtypes of human breast cancer. Since all of these models are on an FVB/NJ inbred mouse background, we bred MMTV-PyMT mice to two pairs of closely related mouse strains that have significantly different inherited metastatic susceptibilities (C57BL/6J and C57BL/10J or MOLF/EiJ and CAST/EiJ, low and high metastatic, respectively) in order to more closely replicate human population diversity. To control for potential contamination of the pulmonary metastases by adjacent tissues, normal lung and metastasis-free tumor-conditioned lung tissue were also profiled. Additionally, we collected tissues from frequently used syngeneic allograft models of mammary tumor metastases. Specifically, matched primary and metastatic tumors from orthotopic injection, tail vein injection, and intracardiac injection models of metastatic breast cancer (4T1, 6DT1, Mvt1, and M6) were included in the analysis. Together this transcriptional analysis profiled 235 RNA samples isolated from 107 animals (Supplemental Table 1). Unsupervised principle component analysis (PCA) of RNA-seq data revealed that the samples clustered into three groups consisting almost exclusively of either autochthonous GEMM-derived samples or allograft samples, with the third group populated by M6 allograft and C3(1)-TAg samples which model basal-like gene expression (Figure 1A)^18^. Metastatic and primary tumors were relatively evenly distributed within each of the clusters, consistent with previous studies that found a high degree of transcriptional similarity between primary and secondary breast cancer samples^19^ (Figure 1B).

**Figure 1.**
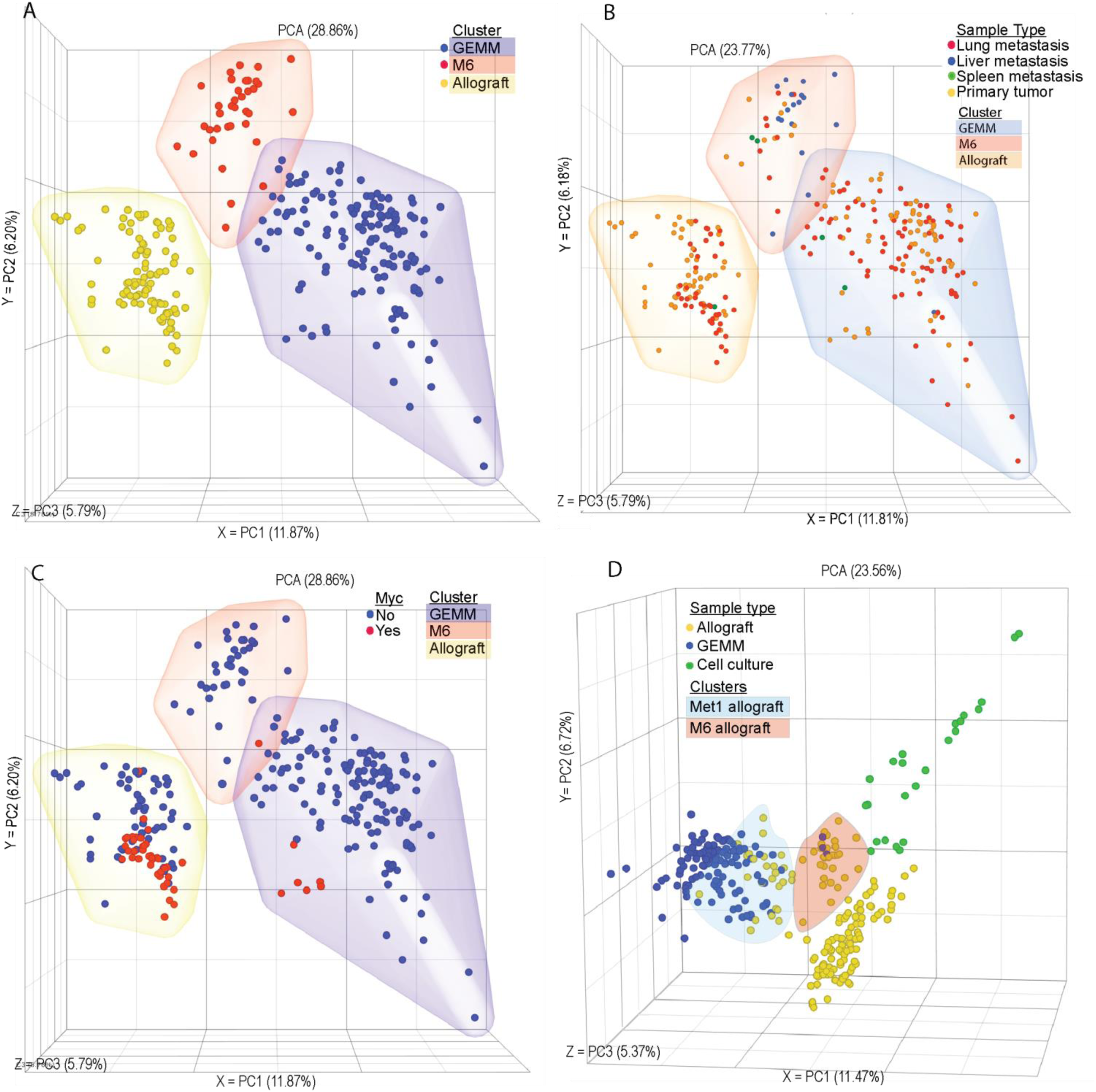
Model has a stronger influence over the tumor transcriptome than driver or site of origin. Unsupervised PCA plot showing RNA-seq transcriptomic analysis from matched primary tumor and metastatic nodules. Each point represents one tissue sample from (A) GEMM (blue), M6 (red), or allograft (yellow) models. (B) Lung metastasis (red), liver metastasis (blue), spleen metastasis (green), primary tumor (yellow). (C) Sample from a Myc-driven tumor model (red), from a non-Myc-driven tumor model (blue) that is a GEMM model (blue cloud), M6 (red cloud), or allograft (yellow cloud). (D) Allograft samples (blue), GEMM samples (yellow), or cell culture samples (red), with Met1 allograft samples (green cloud) and M6 samples (yellow cloud) highlighted.

Allograft models are widely used in the study of metastasis as a model of spontaneous disease, however unsupervised PCA analysis of RNA-seq data revealed that samples clustered by model rather than driver or origin. Indeed, despite being derived from the MMTV-Myc model, both primary and metastatic samples from the 6DT1 and Mvt1 cell line allograft models displayed significant separation from the autochthonous MMTV-Myc tumors, suggesting that *in vitro* growth may result in a permanent transcriptional reprogramming in these cells (Figure 1C). To further explore the potential effect of *in vitro* culture on allograft transcriptional programming, primary tumors from an additional 11 metastatic and non-metastatic mammary tumor allograft models were profiled, in addition to *in vitro* cultures from 7 cell lines (Supplemental Table 1). Allograft tumors from the MMTV-PyMT-derived MET1 cell line clustered with the autochthonous MMTV-PyMT samples, and those from the C3(1)-TAg-derived M6 cell line clustered with the C3(1)-TAg samples between the basal and luminal autochthonous tumor samples (Figure 1D). The remaining nine allograft tumors clustered with the original allograft cluster, regardless of oncogenic driver or genetic background. Finally, *in vitro* samples formed a fourth additional cluster adjacent to the allograft samples (Figure 1D). Together these data reveal that models of metastatic breast cancer differ significantly based on the origin of the transformed cells, and that *in vitro* culturing of metastatic samples permanently alters their transcriptional profile.

To better understand the transcriptional differences between the allograft and GEMM models, variances in the transcriptional profiles were examined. Gene set enrichment analysis of differentially expressed genes between the GEMM and allograft clusters revealed significant differences in immune regulation- and cell motility-related genes (Supplemental Table 2). Analysis of the differentially expressed genes using the ImmQuant software package suggested that the autochthonous GEMM tumors and metastases had decreased levels of infiltrating macrophages, dendritic cells, natural killer cells, and T cells compared to the allograft tumors (Supplemental Figure 1)^20^. Moreover, the GEMM cluster had lower expression of genes associated with the mesenchymal phenotype (e.g., *Vim, Zeb1*), compared to the allograft tumor cluster, and relatively high expression of epithelial markers such as *Epcam, Cldn1*, and *Cldn2* (Figure 2A). Similarly, the MSGE differences between GEMM and allograft models resided mostly in developmental, immune signaling, and EMT pathways (Figure 2B, Supplemental Table 3). This suggests that key differences exist between GEMM and allograft breast cancer metastasis models that involve contributions from both the microenvironment and cell plasticity.

**Figure 2.**
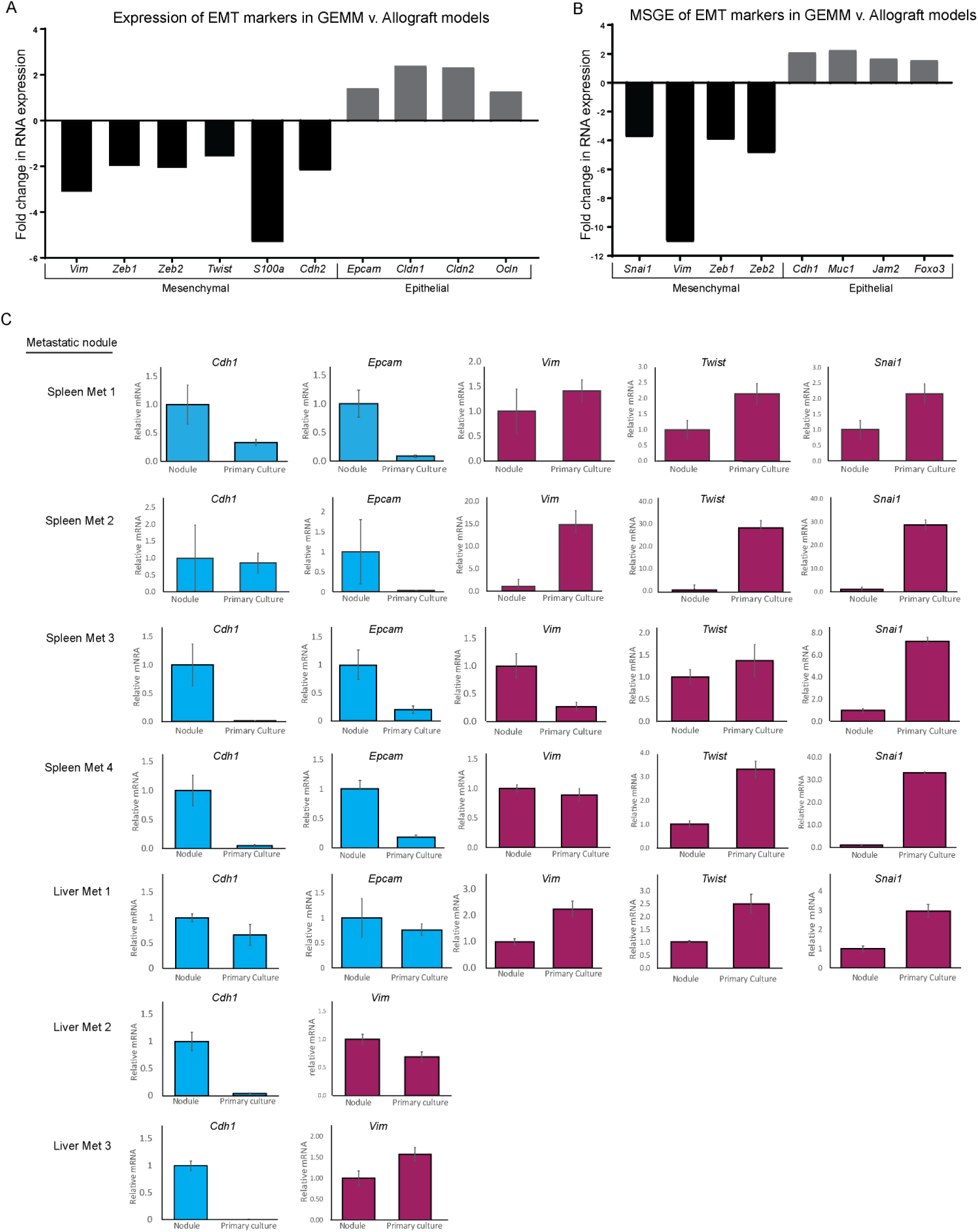
Model is a stronger determinant of EMT signature than tissue disease stage. RNA-seq data showing the fold change in expression of EMT markers in (A) All GEMM derived tissues compared to all allograft-derived tissues, or (B) Metastasis-specific gene expression from GEMM models compared to allograft models. (C) qRT-PCR data showing epithelial (blue) or mesenchymal (red) EMT marker expression in spleen metastases and liver metastases that were subsequently grown in primary culture.

To further explore the possibility that *in vitro* cell culture may promote epithelial to mesenchymal transition (EMT), a gene expression program widely associated with tumor progression and metastatic spread, we measured expression of EMT markers in tissue from an MMTV-Myc mouse harboring lung, liver, and spleen metastases. The spleen and liver metastases collected from this mouse were cut in half and flash frozen, the remaining tissue was then used to create primary cultures which were later injected orthotopically back into ten wild type mice per culture. qRT-PCR analysis was performed on RNA isolated from the original metastases, subsequent primary cultures, and two of the ten injected orthotopic primary tumors, which were each sampled three times to account for primary tumor heterogeneity (Supplemental Figure 2). Pulmonary micro-metastases were observed in 76% of animals (53 out of 70), while only 2.8% (2 or 70) and 1.4% (1 or 70) had liver or spleen metastases, respectively. As observed in our RNA-seq analysis of well-established allograft and GEMM models, culturing of the MMTV-Myc GEMM-derived liver and spleen metastatic cells *in vitro* for < 5 passages imparted a more mesenchymal-like gene expression pattern and reduced epithelial gene expression (Figure 2C). This change in gene expression was variably maintained once cells were implanted orthotopically *in vivo* (Supplemental Figure 3). Taken together, this data suggests that *in vitro* culturing techniques induce a more mesenchymal and lung-tropic phenotype that variably persists and does not consistently or accurately model the plasticity and tropism of metastatic cells as they originally existed *in vivo*.

### Orthotopically derived and tail vein-injected metastases differ in a cell line-dependent manner

The preparation of a premetastatic niche at the secondary site is now widely accepted as a key step in the metastatic process^21–23^. Therefore, we examined if metastasis assays performed by tail vein injection, in the absence of a primary tumor and premetastatic niche conditioning, produced metastases with different transcriptional profiles than those arising from orthotopically injected cells. To address this, the MSGE of lung metastases derived from syngeneic tail vein injection or orthotopic injection of three metastatic mouse cell lines (4T1, Mvt1, and 6DT1) was established (Supplemental Figure 4). The orthotopic injection MSGE was then compared to tail vein injection MSGE for each cell line individually (Figure 3A).

**Figure 3.**
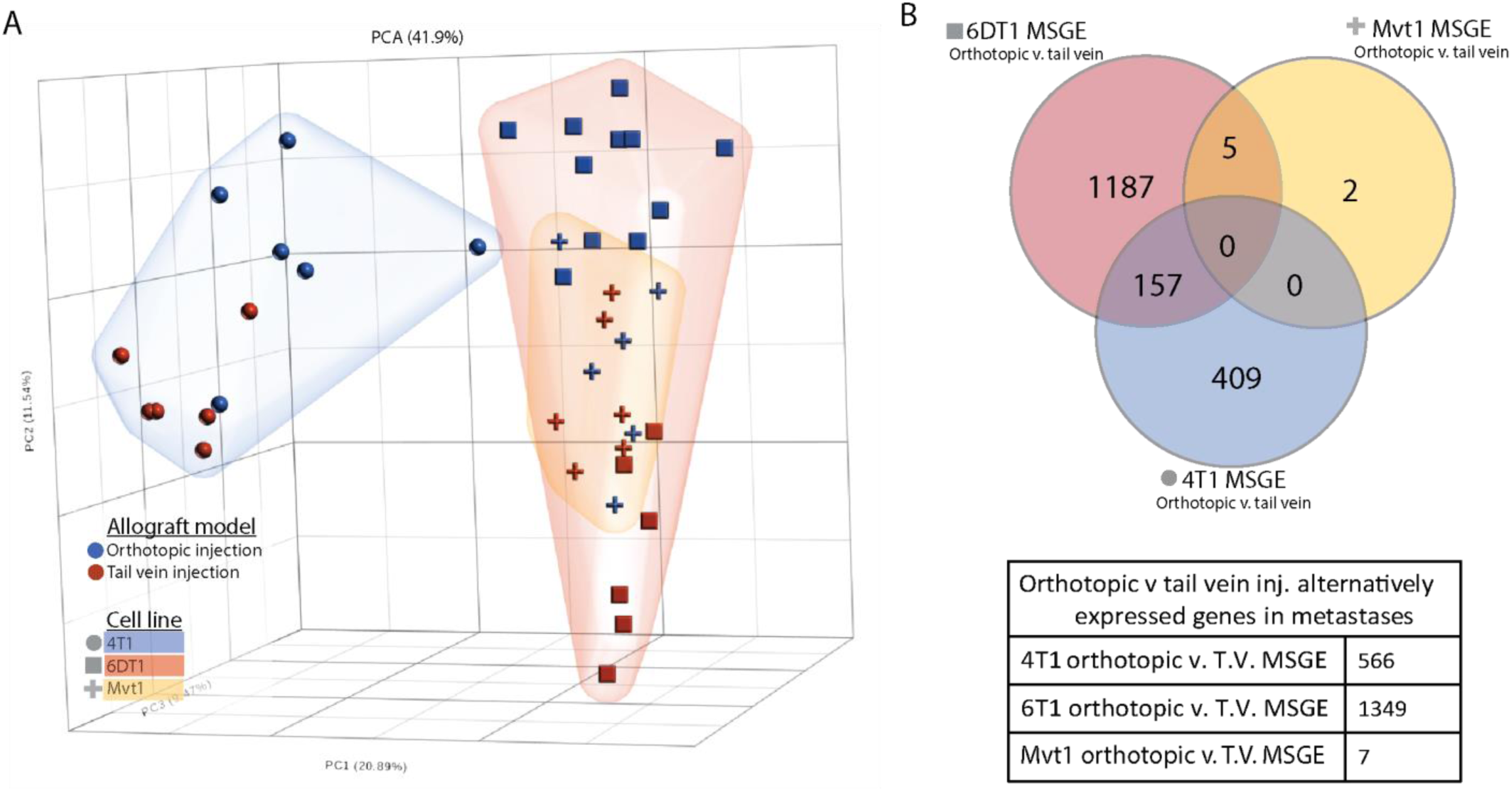
Metastases derived from orthotopic and tail vein injection models differ in a cell line-dependent manner. (A) PCA analysis plot showing relative difference between the transcriptomes of tail vein-derived lung metastases (red points) and orthotopic injection model-derived lung metastasis (blue points), with 4T1 cells (blue cloud, circles), 6DT1 cells (red cloud, squares), and Mvt1 cells (yellow cloud, crosses). (B) Venn diagram showing the overlapping orthotopic-specific MSGE from the 4T1 (blue), 6DT1 (red), and Mvt1 (yellow) allograft models.

Interestingly, this analysis revealed that the differences in metastatic gene expression between models was cell line-dependent. Metastases derived from orthotopic or tail vein injection of Mvt1 cells differed only in the expression of 7 genes, but metastases derived from 4T1 cell injections differed in the expression of 566 genes between injection models, and 6DT1 derived metastases differed by 1349 genes (Figure 3B; Supplemental Table 4). There were no genes commonly altered by injection model across all three cell lines (Figure 3B). However, differentially expressed genes specific to each cell line were assessed by Ingenuity Pathway Analysis (IPA) (Table 1 and Supplemental Table 5), which revealed that the gene expression activated specifically in orthotopically derived metastases populated T cell activation-related pathways when compared to tail vein-derived metastases. This suggests that metastatic tumors derived from these two models may undergo different interactions with the immune compartment at the metastatic site and differ significantly in gene expression in a cell line-dependent manner.

**Table 1.** Ingenuity canonical pathways of orthotopic allograft model MSGE. Ingenuity Pathway Analysis results showing the pathways populated by orthotopic-specific MSGE of 4T1, 6DT1, and Mvt1. Significance of enrichment for this pathway in each the gene set is indicated by the −log(p-value) where >1.3 is significant.

| Table 1: Ingenuity canonical pathways of orthotopic allograft model MSGE |  |  |  |  |  |
| --- | --- | --- | --- | --- | --- |
| 6DT1 | -log(p-value) | 4T1 | -log(p-value) | Mvt1 | -log(p-value) |
| Th1 and Th2 Activation Pathway | 17.5 | Agranulocyte Adhesion and Diapedesis | 9.76 | Autophagy | 2.06 |
| Th2 Pathway | 13.2 | Granulocyte Adhesion and Diapedesis | 8.76 | Th1 Pathway | 1.75 |
| Th1 Pathway | 11.5 | Primary Immunodeficiency Signaling | 7.92 | Th2 Pathway | 1.71 |
| Granulocyte Adhesion and Diapedesis | 10.1 | T Helper Cell Differentiation | 7.83 | Th1 and Th2 Activation Pathway | 1.61 |
| T Helper Cell Differentiation | 9.12 | Th1 and Th2 Activation Pathway | 7.76 | Sirtuin Signaling Pathway | 1.38 |
| Altered T Cell and B Cell Signaling in Rheumatoid Arthritis | 8.16 | Th2 Pathway | 6.99 |  |  |
| Agranulocyte Adhesion and Diapedesis | 7.6 | Th1 Pathway | 6.13 |  |  |

### Nicotine and Sildenafil processing are targetable, metastasis-specific transcriptional programs

As metastatic nodules in human patients are often resistant to treatments that the primary tumor was sensitive to, there is a clear need for metastasis-specific therapies. We next examined targetable transcriptional programs common to spontaneous metastases in both allograft and GEMM models. First, the combined MSGE for spontaneous allograft models was determined for each cell line. MSGE common across 6DT1, 4T1, and Mvt1 orthotopically injected cell lines was then compared to MSGE of GEMMs (Supplemental Figure 5). 360 genes were commonly regulated in metastases from both allograft and autochthonous models of spontaneous metastasis. IPA of this gene set analysis revealed several signaling, motility, and metabolic pathways in addition to several targetable pathways including “Cellular Effects of Sildenafil Citrate (Viagra)” and “Nicotine Degradation” (Table 2, Figure 4A and 4B, Supplemental Table 6). Interrogation of these pathways as regulators of breast cancer metastasis was performed by spontaneous metastasis assays using orthotopic injection of metastatic breast cancer 6DT1 cells in the presence of sildenafil citrate or nicotine.

**Table 2.** Ingenuity canonical pathways of MSGE common to spontaneous models of metastasis. Ingenuity Pathway Analysis results showing the pathways populated by MSGE common to both spontaneous allograft and GEMM models. Significance of enrichment for this pathway in each the gene set is indicated by the −log(p-value) where >1.3 is significant.

| Table 2: Ingenuity canonical pathways of MSGE common to spontaneous models of metastasis | -log(p-value) |
| --- | --- |
| Nicotine Degradation II | 6.98 |
| Estrogen Biosynthesis | 5.23 |
| Acetone Degradation I (to Methylglyoxal) | 5.13 |
| Bupropion Degradation | 4.4 |
| Agranulocyte Adhesion and Diapedesis | 3.94 |
| Nicotine Degradation III | 3.66 |
| Hepatic Fibrosis / Hepatic Stellate Cell Activation | 3.35 |
| Atherosclerosis Signaling | 3.23 |
| Intrinsic Prothrombin Activation Pathway | 3.04 |
| Cellular Effects of Sildenafil (Viagra) | 2.91 |
| Granulocyte Adhesion and Diapedesis | 2.88 |

**Figure 4.**
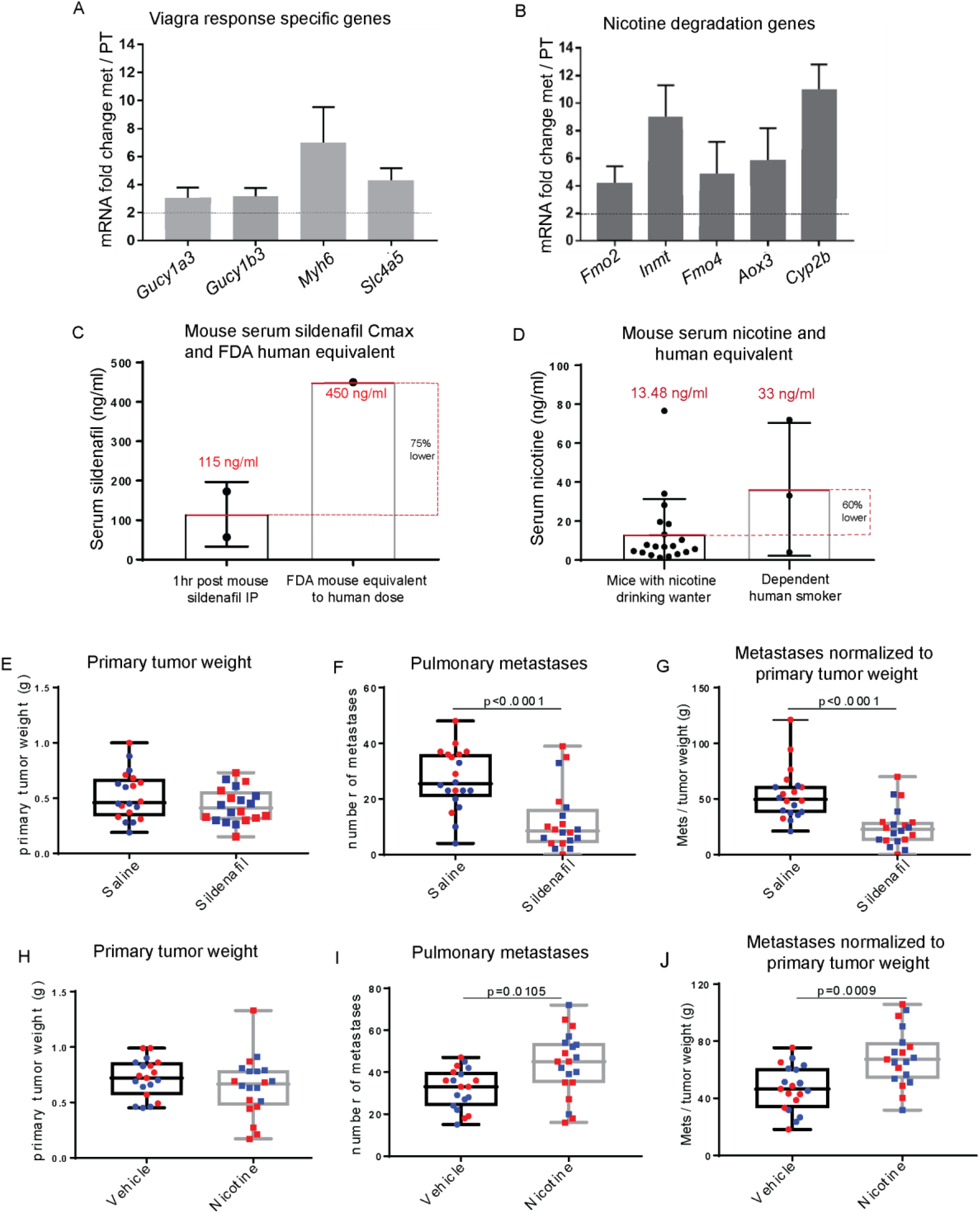
Common MSGE of several models of spontaneous metastasis present targetable metastasis-specific pathways. RNA-seq data showing expression of factors from the IPA sildenafil response pathway (A) and Nicotine degradation pathways (B) in metastatic v. primary tumor tissue. (C) Max Serum (Cmax) of sildenafil 1hr post IP injection of 10 mg/kg sildenafil (each point corresponds to one mouse) compared to the FDA mouse equivalent sildenafil serum levels of a human dose (D) Serum levels of nicotine achieved in this study following constant access to nicotine 100 μg/ml drinking water (each point corresponds to one mouse) compared to the average nicotine serum levels of dependent human smokers (range 4 – 72 ng/ul, average 33 ng/μl^27^). (E) Primary tumor weight (g), (F) lung nodule counts, and (E) lung nodule counts normalized to paired primary tumor weight at end point following syngeneic orthotopic injection of 1×10^5^ 6DT1 cells and treatment with saline (circles) or sildenafil (10 mg/kg) (squares). (H) Primary tumor weight (g), (I) lung nodule counts, and (J) lung nodule counts normalized to paired primary tumor weight at end point following syngeneic orthotopic injection of 1×10^5^ 6DT1 cells and access to nicotine or vehicle water (E-J). Two independent experiments shown (red and blue).

To determine if sildenafil citrate treatment could modify breast cancer metastasis, FVB/NJ mice bearing 6DT1 orthografts were treated with a daily low dose of sildenafil citrate by intraperitoneal injection starting 7 days post-injection of cells and lasting 21 days until endpoint. Pharmacokinetic/pharmacodynamic (PK/PD) analysis of serum collected at several time points following sildenafil citrate injection indicated that a Cmax of 115 ng/ml was achieved at 1hr post sildenafil IP, a serum level equivalent to 25% of typical human dose according to FDA guidelines for human and mouse equivalent dosing (Figure 4C, Supplemental Table 7)^24–26^. While treatment with sildenafil did not significantly alter primary tumor growth it did significantly reduce the number of surface metastases on the lung (p < 0.0001) (Figure 4E-G), indicating that sildenafil processing is an integral, metastasis-specific pathway necessary for efficient metastatic spread.

To determine if cellular nicotine degradation pathway activity is a modifier of breast cancer metastasis, FVB/NJ mice bearing 6DT1 orthografts were administered nicotine in their drinking water starting 7 days post-injection of cells and lasting 21 days until endpoint. PK/PD analysis of nicotine and its metabolite cotinine in serum collected throughout a 4-week course of nicotine treatment indicated that mouse nicotine serum levels averaged approximately 13.48 ng/ml, which is 60% lower than that of a dependent smoker (Figure 4D, Supplemental Figure 8)^27–30^. Treatment with nicotine did not alter primary tumor growth but did significantly increase the number of surface metastases on the lung (p < 0.001) (Figure 4H-J). Again, this data suggests that the activation of nicotine degradation gene expression pathways is a metastasis-specific function resulting in more efficient metastatic spread.

## Discussion

A major boundary to progress in the development of metastasis-specific drugs is a lack of available human tissue for study, forcing researchers to rely heavily on pre-clinical mouse models. Mouse models of metastatic breast cancer are widely used, and a large number of models exist to interrogate oncogenic drivers, genetic backgrounds, and stages of the metastatic cascade^31^. As no single model likely captures the full complexity of metastatic breast cancer observed in humans, we hypothesized that gene expression observed in metastases from each model may exhibit both model-specific as well as common and essential metastatic programs. Therefore, to advance the development of metastasis-specific interventions, we interrogated the transcriptomes of metastatic tissues isolated from a wide variety of pre-clinical mouse models of metastatic breast cancer.

We performed RNA sequencing on paired metastatic and primary tumors from three allograft models using five mouse mammary cancer cell lines, along with four GEMMs crossed to five different strain backgrounds, which together featured more than six clinically relevant oncogenic drivers. Here we have shown that important differences exist in the gene expression profiles of metastases generated by different models, and that the model may play a larger role in shaping the metastatic transcriptome than tissue type or oncogenic driver. Despite these differences, models of spontaneous metastasis do exhibit common metastasis-specific gene expression patterns that can be readily targeted. Using the 6DT1 orthotopic injection allograft model as a proof of concept, we show that a low dose of sildenafil citrate significantly reduced pulmonary metastasis, whereas a low dose of nicotine significantly promoted metastasis. These data, along with our continued characterization of additional GEMM and allograft models of metastatic breast cancer, will improve selection of pre-clinical models for research and better inform the interpretation of subsequent results. Additionally, we anticipate that the combined analysis of multiple models will uncover further metastasis-essential gene expression pathways for the development of metastasis-specific interventions.

EMT is postulated to be a key mechanism by which tumor cells can acquire an invasive and metastatic phenotype. However, most of the evidence for the existence of EMT in cancer is derived from *in vitro* experimental systems and cancer cell lines^32^. In fact, work by Trimboli et al. on EMT in PyMT, Neu, and Myc GEMMs shows that EMT-like gene expression signatures are only observed for the Myc GEMM, despite the high metastatic propensity of the PyMT model^33^. Similarly, our data suggest that the EMT phenomenon is not predominant in PT or metastatic tissues collected from breast cancer GEMM models, as the samples analyzed in this study overwhelmingly exhibit more epithelial-like gene expression profile compared to cell-line-derived allograft tissue samples. Interestingly, by PCA analysis, MMTV-Myc tissue samples cluster with the remaining GEMM samples but are located on the periphery of this cluster closest to the allograft cluster, potentially signifying this model’s propensity for an EMT-like phenotype. Additionally, our additional analysis of metastatic tissue grown in primary culture further supports the observation that *in vitro* culturing of mouse mammary cancer cells may impart a more mesenchymal-like gene expression signature. Together our data suggest that EMT may be a marker of model rather than metastatic potential when working with mouse models of metastatic breast cancer.

Despite the necessity of mouse models for the study of metastasis, little is known about the differences between the many models so frequently employed. Indeed, it is unclear how different model-specific pressures, such as the need to intravasate (orthotopic injection allograft models and GEMMs) or not (intravenous injection allograft models), may impact the biology of the resulting metastatic nodules. Our analysis of metastases isolated from either orthotopic injection or tail vein injection of the same cell line revealed key model-specific differences in MSGE, the extent of which was cell line-dependent. Interestingly, the variance in gene expression observed between these routes of implantation implicated immune interaction as a key distinguishing factor. It is now widely accepted that factors released from the primary tumor prepare the secondary organ, and altered immune infiltration is a major aspect of this niche formation^22,23,34–36^. Based on our data, we hypothesize that the pre-metastatic niche may be key in shaping the MSGE of some cell lines when used in allograft models. Again, this result suggests that complex model-specific variances likely play a large role in shaping the biology of the resultant metastatic tumors.

Despite the significant transcriptional differences, we have also identified common gene expression patterns in metastases derived from several models of spontaneous metastasis. Our data revealed an upregulation of factors responsible for the cellular effects of sildenafil citrate (Viagra), which is an inhibitor of phosphodiesterase type 5 (PDE5) and ultimately a regulator of cellular cGMP^37^. Clinically, sildenafil is administered almost exclusively to men, and therefore no data is available to interrogate the effects of sildenafil on breast cancer patient outcomes. However, the target of sildenafil, PDE5, has been identified as a tumor biomarker in several human cancers^38–41^, and work with human cancer cell lines has revealed that expression and activation of PDE5 stimulates invasive, migratory, and cancer stem cell phenotypes ^41,42^. Here we show that a low dose of sildenafil, administered by intraperitoneal injection, significantly reduced spontaneous metastasis of the 6DT1 mouse mammary cancer cell line. While our transcriptomics data and research by others would suggest this reduction in metastatic potential is likely tumor cell-intrinsic^41,42^, we cannot rule out the possibility that sildenafil may also induce a systemic contribution to this phenotype. Furthermore, this discovery forces us to consider at which stage in the disease this compound may be effective, and if there are additional FDA-approved compounds that may influence the progression of metastatic cancers.

In addition to the cellular effect of sildenafil, we identified an increase in factors responsible for nicotine degradation across metastatic tissues collected from spontaneous metastasis models. Since the mice were not exposed to nicotine, we hypothesize that metabolites generated by this pathway may promote metastasis. Administration of nicotine through drinking water significantly increased the rate of metastasis in an orthotopic injection model using the 6DT1 cell line. Again, while our transcriptomics data suggests this response is likely tumor cell-intrinsic, we cannot rule out the possibility that nicotine ingestion may also induce systemic contributions to this highly metastatic phenotype. More work will be necessary to understand the cellular processing of nicotine and subsequent effects on tumor cell behavior. Patient data show that breast cancer patients who are current or former smokers have a higher incidence of disease progression and mortality^43^. While current human data sets cannot distinguish the effect of nicotine from tobacco, our data suggest that exposure to nicotine alone may be pro-metastatic, implicating tobacco-free nicotine products as potentially harmful to breast cancer patients. Taken together, these data suggest that the sildenafil response and nicotine metabolism pathways may be useful and clinically actionable targets for adjuvant or neoadjuvant therapy.

In this study, we crossed the MMTV-PyMT GEMM to five mouse strains to better account for diversity in the human population. Despite the large number of mice included in this study, we do not currently have enough statistical power to draw conclusions about the impact of strain on MSGE. Our ongoing work endeavors to assess the metastatic transcriptome in additional strains while also increasing the number of samples derived from each strain in order to identify strain-specific features and potential stromal programing of the metastatic transcriptome. This will complement our previous findings that genetic background influences metastatic potential in the MMTV-PyMT model^44–47^. Furthermore, we plan to continue tissue collection and MSGE analysis of additional MMTV-Her2, Myc, and C3(1)-TAg animals to compare MSGE between GEMMs with varying oncogenic drivers. This effort will provide valuable insight into the different vulnerabilities of metastases from patients of different subtypes. Finally, while our data targeting metastasis-specific pathways with sildenafil citrate and nicotine is striking, additional work will be required to determined how these compounds influence the metastatic cascade. We are particularly interested to address whether sildenafil can reduce the size of already established metastases, and if nicotine can promote the dissemination of non-metastatic tumors.

In summary, by performing a survey of the MSGE profiles of macrometastases from several allograft models and GEMMs, we have discovered key differences that exist between models as well as core metastasis-specific programs that can be readily targeted. By using the strategy outlined here, the continued characterization of preclinical mouse models of metastasis will provide further insight into the associations between clinical subtypes, primary tumor drivers, and MSGE programs, ultimately creating new opportunities for the generation of metastasis-targeted therapies.

## Materials and Methods

### Ethics statement

The research described in this study was performed under the Animal Study Protocol LCBG-004 and LPG-002, approved by the National Cancer Institute (NCI) Animal Use and Care Committee. Animal euthanasia was performed by cervical dislocation after anesthesia by Avertin.

### Genetically engineered mouse models

FVB/N-Tg(MMTV-PyVT)634Mul/J and FVB/N-Tg(MMTVneu)202Mul/J male mice were obtained from The Jackson Laboratory. FVB/N-Tg(C3(1)-TAg)^48^ and FVB/N-Tg(MMTV-Myc) were a generous gift from Dr. Jeffrey Green (NCI, Bethesda, MD).

Male PyMT mice were crossed with female wild type FVB/NJ, MOLF/EiJ, CAST/EiJ, C57BL/6J, and C57BL/10J mice also obtained from The Jackson Laboratory. Male Her2, Myc, and C3(1)-TAg mice were crossed with female wild type FVB/NJ mice. All female F1 progeny were genotyped by the Laboratory of Cancer Biology and Genetics genotype core for the PyMT or Her2 gene and grown until humane endpoint. Primary tumor, metastatic nodules, and normal (tail) tissue were isolated immediately following euthanasia and snap-frozen in liquid nitrogen. Tissue samples were then stored at −80°C.

### Cell culture

The mouse mammary carcinoma cell lines 6DT1, 4T1, Mvt1, and MET1 were a generous gift from Dr. Lalage Wakefield (NCI, Bethesda, MD). The mouse mammary carcinoma cell line M6 was a generous gift from Dr. Jeffrey Green (NCI, Bethesda, MD). All cell lines were cultured in Dulbecco’s Modified Eagle Medium (DMEM), supplemented with 10% Fetal Bovine Serum (FBS), 1% Penicillin and Streptomycin (P/S), and 1% L-Glutamine (Gibco), and maintained at 37°C with 5% CO_2_.

### *In vivo* metastasis

Female virgin FVB/NJ or BALB/cJ mice were obtained from The Jackson Laboratory. Two days prior to *in vivo* experiments, cells were plated at 1 × 10^6^ cells/condition into T-75 flasks (Corning) in non-selective DMEM. A total of 100,000 cells in 100 µl of PBS were injected per mouse into the fourth mammary fat pad (orthotopic injection), tail vein (tail vein injection), or left ventricle (intracardiac injection). The mice were euthanized between 28–30 days post-injection. Primary tumors were resected, weighed, and lung metastases counted. Primary tumor, metastatic nodules, and normal (tail) tissue were isolated immediately following euthanasia and snap-frozen in liquid nitrogen. Tissue samples were then stored at −80°C. For orthotopic injection resection models, the primary tumors were resected at 2 weeks following initial injection. and mice were euthanized between 42-45 days post-injection. Primary tumors from syngeneic orthotopic injection of r3T, D2A1, E0771, EMT6, F311, HRM-1, and TS/A-E1 were a generous gift from Dr. Lalage Wakefield (NCI, Bethesda, MD).

#### Syngeneic models

FVB/NJ - 6DT1, Mvt1, Met1, M6C. BALB/cJ – 4T1, 4T07, 67NR

#### Sildenafil citrate treatment

6DT1 orthotopic injection was performed as described above. Dosing with 10 mg/kg sildenafil citrate was initiated on day 7 post-injection and was repeated daily until day 28, the study end-point. Sildenafil citrate salt (Sigma-Aldrich) was resuspended in sterile PBS (Life Technologies) to create a 5 mg/ml stock solution, which was stored at −20°C. Each mouse was weighed daily before dosing and 10 mg/kg dose was calculated and then administered by diluting the sildenafil citrate stock solution with sterile PBS to a final volume of 250 µl. Control mice were weighed and administered 250 µl of PBS. Mice were administered sildenafil citrate by IP injection of 250 µl using a 28-gauge needle (Andwin Scientific).

#### Nicotine administration

6DT1 orthotopic injection was performed as described above. On day 7 following injection mice were given drinking water with nicotine or vehicle alone. (-)-Nicotine (Sigma-Aldrich) was diluted in 100% ethanol to create a stock solution with a final concentration of 100 mg/ml. 250 µl of the nicotine stock solution, or 250 µl of ethanol was added to a 250 ml of mouse drinking water for a final concentration of 100 µg/ml. The water was replaced weekly and the nicotine stock solution was made fresh each week.

### Serum isolation

Following mouse anesthesia, 500-1000 µl of blood was removed by cardiac puncture using a 25-gauge needle. Blood was stored on ice with 10 µl of EDTA (0.5M, pH 8) and then centrifuged at 2,000 x g for 15 minutes in a refrigerated centrifuge. The plasma supernatant was then immediately removed and placed into a new tube and stored at −20°C.

#### Sildenafil citrate trial serum collection

Serum was collected from tumor bearing mice 1, 4, 6, 8, 18, and 24 hours after the final sildenafil citrate or PBS control IP injection was performed on day 28 (see sildenafil citrate treatment regimen above).

#### Nicotine trial serum collection

20 mice were administered 100 µg/ml nicotine in the drinking water, and 5 mice were administered vehicle control drinking water. Serum was collected from 5 mice in the nicotine group once a week for 4 weeks. Serum was collected from the 5 control mice after the final 4^th^ week. Water was replaced weekly.

### LC-MS/MS drug-serum level assessment

Nicotine and cotinine plasma concentrations were simultaneously measured using a validated LC-MS/MS assay, with a lower limit of quantitation of 1 ng/mL for both compounds. Briefly, 100 µL of plasma was mixed with 1 ml of dichloromethane containing 2 ng/ml of [2H]4-nicotine as internal standard. The organic layer was isolated, dried under nitrogen, reconstituted with (10/90, v/v) water/methanol, and injected into the LC-MS/MS. The calibration range for both nicotine and cotinine were 1-500 ng/ml.

Sildenafil plasma concentrations in mice dosed orally were measured using a validated LC-MS/MS assay. Briefly, 50 µl of plasma sample was mixed with 5-fold volume of methanol containing 100 ng/ml 2[H]8-sildenafil (as internal standard) to precipitate plasma proteins. The mixture was vortexed and centrifuged, and 200 µl of the resulting supernatant was transferred to a clean 96-well sample plate for measurement. A 10-µL aliquot of sample was injected onto a Kinetex C18 column (50 × 2.1 mm, 1.3 µm; Phenomenex) for chromatographic separation followed by multiple reaction monitoring detection by tandem mass spectrometry (MS/MS) by the mass transition of m/z 475.1 to 283.0 for sildenafil and m/z 482.5 to 283.0 for 2[H]8-sildenafil. Calibration standards of sildenafil were prepared over the range of 2.5 – 2500 ng/ml (in duplicate) with quality control (QC) standards at a low (7.5 ng/ml), mid (400 ng/ml) and high (2000 ng/ml) range, each in quintuplet

### RNA isolation

RNA was isolated from flash-frozen tissue. First the tissue was mechanically dissociated using a tissue grinder while submerged in 1 ml of TriPure (Roche). 200 µl of chloroform (Sigma-Aldrich) was then added and the soluble fraction was isolated by centrifugation at 12,000 rpm for 15 minutes at 4°C. RNA was then precipitated with the addition of 500 µl isopropanol and incubation of the sample at −20°C for 2 hours. Pure RNA was then extracted using the RNA: DNA mini-prep kit (Zymogen) and finally samples were eluted in 100 µl (PT) or 50 µl (metastases) of DEPC water (Quality Biological). RNA was isolated from cell lines using TriPure as described above but following isopropanol precipitated RNA was washed with 75% ethanol (Sigma-Aldrich), and then again with 95% ethanol before being resuspended in 100 µl DEPC water.

### RNA sequencing

RNA quality was tested using the Agilent 2200 TapeStation electrophoresis system, and samples with an RNA integrity number (RIN) score >7 were sent to the Sequencing Facility at Frederick National Laboratory. Preparation of mRNA libraries and mRNA sequencing was performed by the Sequencing Facility using the HiSeq2500 instrument with Illumina TruSeq v4 chemistry.

### RNA sequencing analysis

#### Differential gene expression analysis

RNA-seq reads were aligned to the mouse mm9 genome assembly using TopHat Software, and differentially expressed genes were determined using DESeq2.

#### Gene Set Enrichment Analysis (GSEA)

The hierarchical cluster analysis was applied to the quantile normalized RPKM data of the primary tumor and metastatic samples to obtain two groups. One group had 88% allograft samples and another group had 86% GEMM samples. DESeq2 was then applied to find the differentially expressed genes between the two groups and then used GSEA to find significant cancer Hallmark pathways and Gene Ontology (GO) pathways.

#### Principal Component Analysis

PCA analysis was performed using Partek Flow software (Kanehisa Laboratories). RNA-seq reads were uploaded into Partek Flow and aligned with the mouse mm9 genome assembly. Gene counts were then determined and normalized before performing unsupervised principal component analysis.

#### ImmQuant analysis

DESeq2 fold-change data for those genes differentially regulated in GEMM and allograft models (Supplemental Table 2) were loaded into the ImmQuant Software^20^. The following settings were selected for default deconvolution: Cell-type data - ImmGen 207, signature markers, lineage tree pre-compiled based on the reference data, and calculations were performed using all samples. Default color ranges were also selected for the lineage tree output file.

#### Pathway Analysis

Pathway analysis was performed using Ingenuity Pathway Analysis (IPA) (Qiagen). Differentially regulated gene lists from sample type comparisons using DESeq2 were uploaded into IPA for Core Expression Analysis of expression data. The Ingenuity Knowledge Base was chosen as the reference set of genes, and both direct and indirect relationships were considered. No other analysis parameters were specified, and the default settings were selected.

### qRT-PCR

RNA was isolated from cell lines or flash-frozen tissue as described above was reverse-transcribed using iScript (Bio-Rad). Real-Time PCR was conducted using VeriQuest SYBR Green qPCR Master Mix (Affymetrix). Peptidylprolyl isomerase B (*Ppib*) was used for normalization of expression levels. Expression of mRNA was defined from the threshold cycle, and relative expression levels were calculated using delta delta Ct after normalization with *Ppib*.

### Statistics

Statistical significance between groups in *in vivo* assays was determined using the Mann-Whitney unpaired nonparametric test using Prism (version 5.03, GraphPad Software, La Jolla, CA). Statistical significance between samples in qRT-PCR analysis was determined by an unpaired t test, also using Prism.

### Data availability statement

All sequence data that supports the findings of this study will be deposited in a public repository and the accession codes will be made available prior to publication.

## Supplemental Figures

**Supplemental Figure 1.**
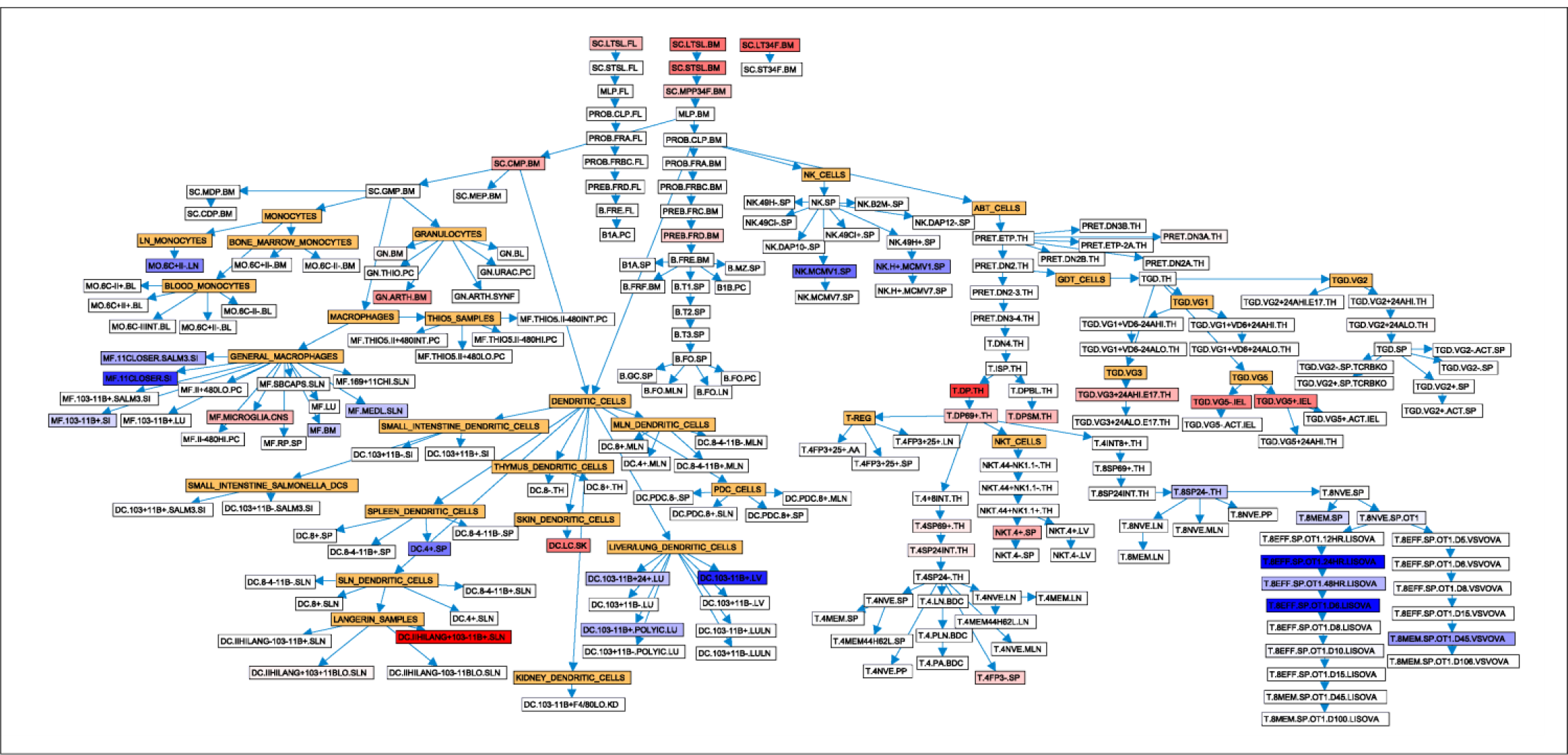
ImmQuant results showing altered immune compartment signaling in GEMM tumor tissue compared to allograft tumor tissue. Hematopoietic lineage nodes are yellow. A predicted increase or decrease in abundance of each immune cell type is indicated by red or blue boxes, respectively.

**Supplemental Figure 2.**
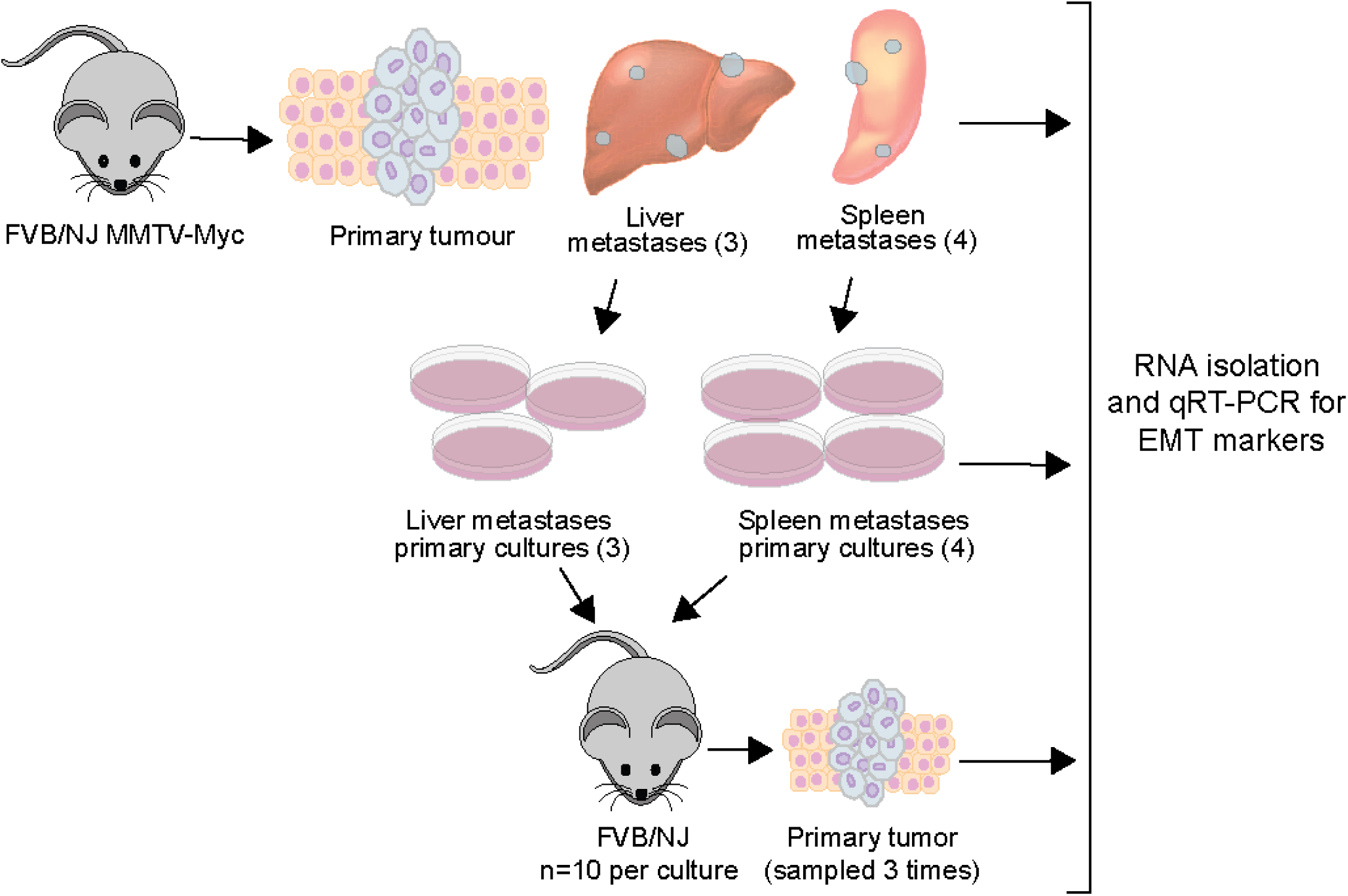
Schematic indicating the workflow of tissue isolation, primary culturing, and implantation back into FVB/NJ mice. Primary and metastatic tumor tissues collected from an FVB/NJ MMTV-Myc mouse with liver and spleen metastases. Those tissues were then used to create primary cultures (Liver:3, Spleen:4), which were then injected orthotopically back into FVB/NJ mice (10 per culture). At each stage, RNA was isolated, and qRT-PCR was performed to measure the expression of EMT markers.

**Supplemental Figure 3.**
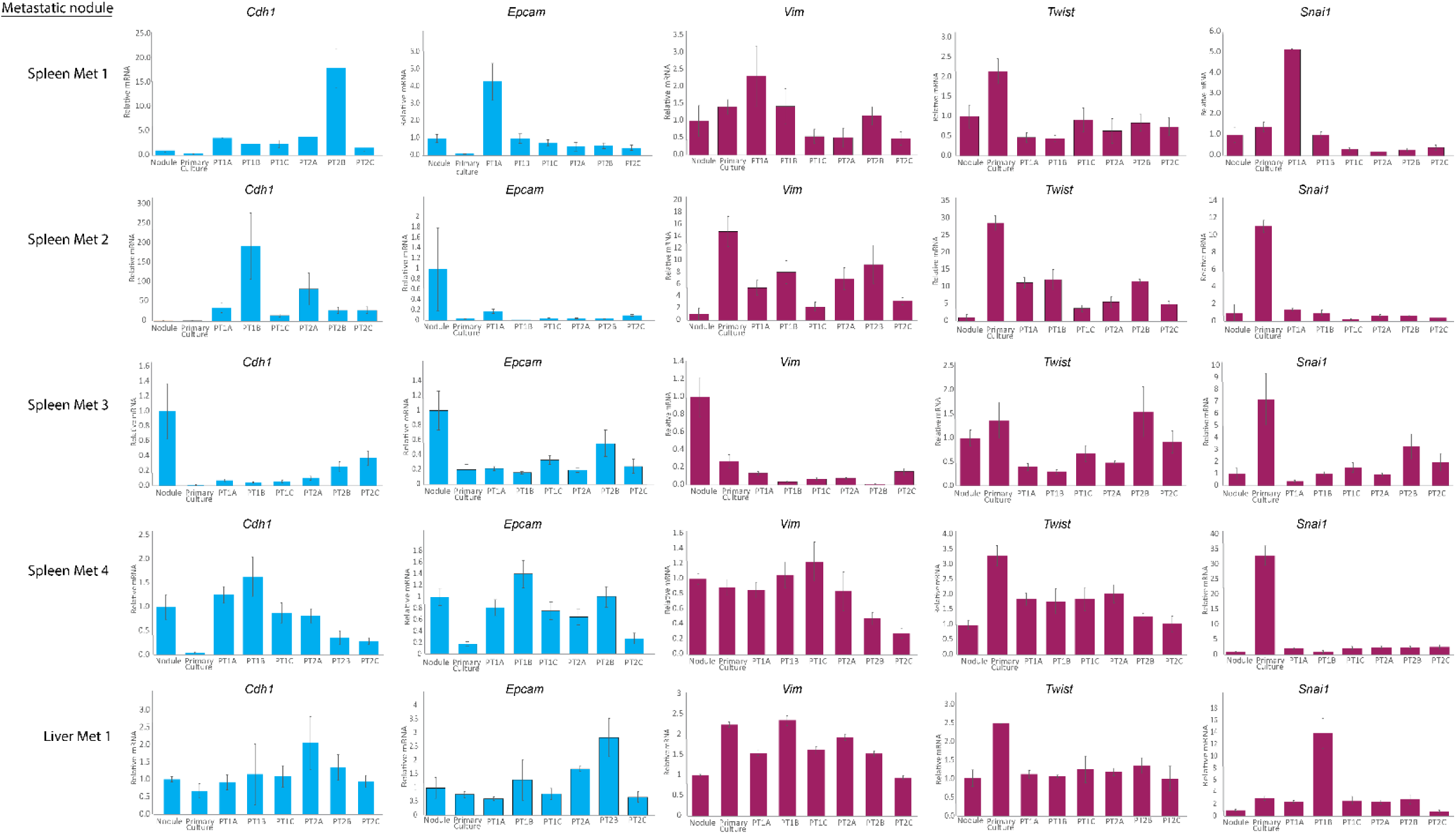
qRT-PCR analysis of EMT markers in RNA isolated from an MMTV-Myc liver, and spleen metastases. Each metastatic nodule was then grown in culture and orthotopically injected back into 10 mice. Shown here is expression data from the original metastatic nodule (Nodule), the primary culture (Primary Culture), and 2 primary tumors sampled 3 times each (PT1A-C, and PT2A-C) from subsequent orthotopic injection of the primary culture. Standard deviation of technical replicates is indicated by bars.

**Supplemental Figure 4.**
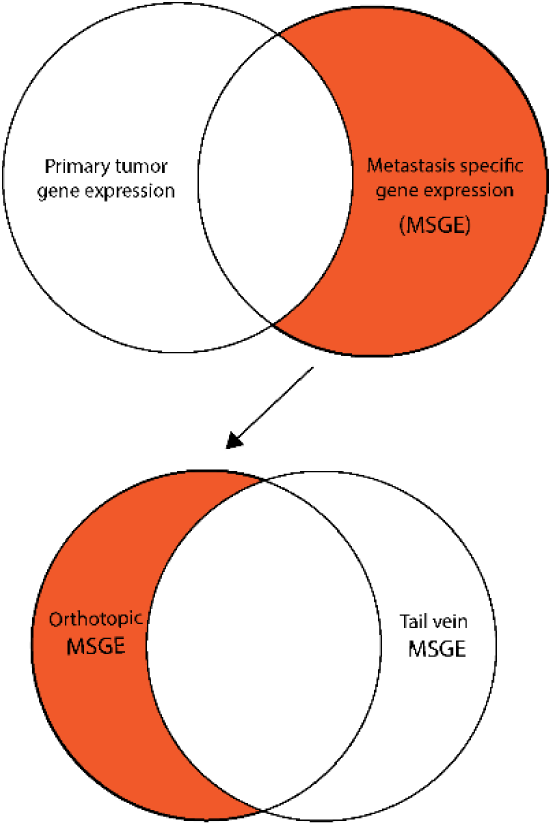
Schematic of basic workflow for identification of orthotopic injection-specific MSGE.

**Supplemental Figure 5.**
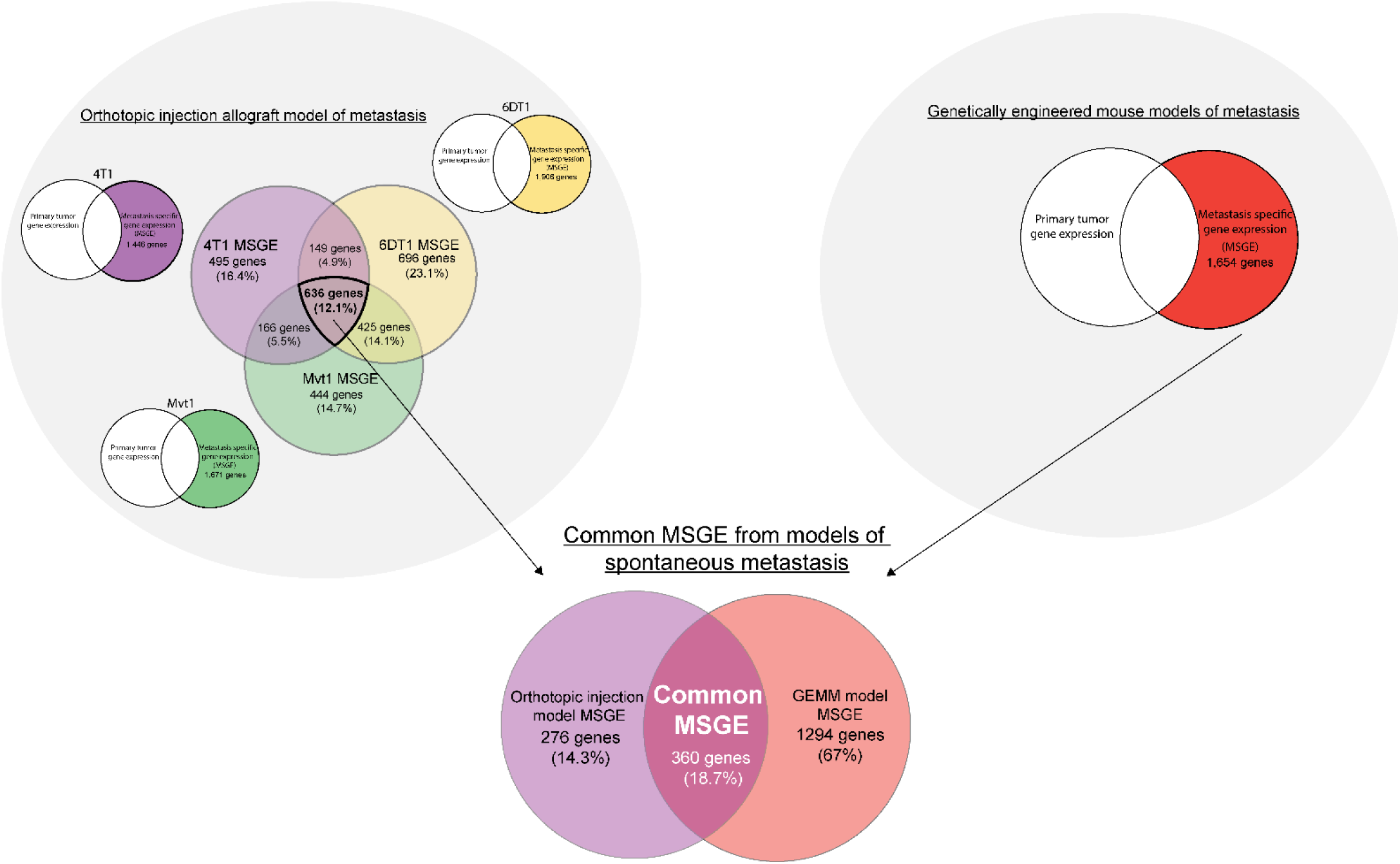
Schematic of basic workflow for the identification of common MSGE in models of spontaneous metastasis.

**Supplemental Figure 6.**
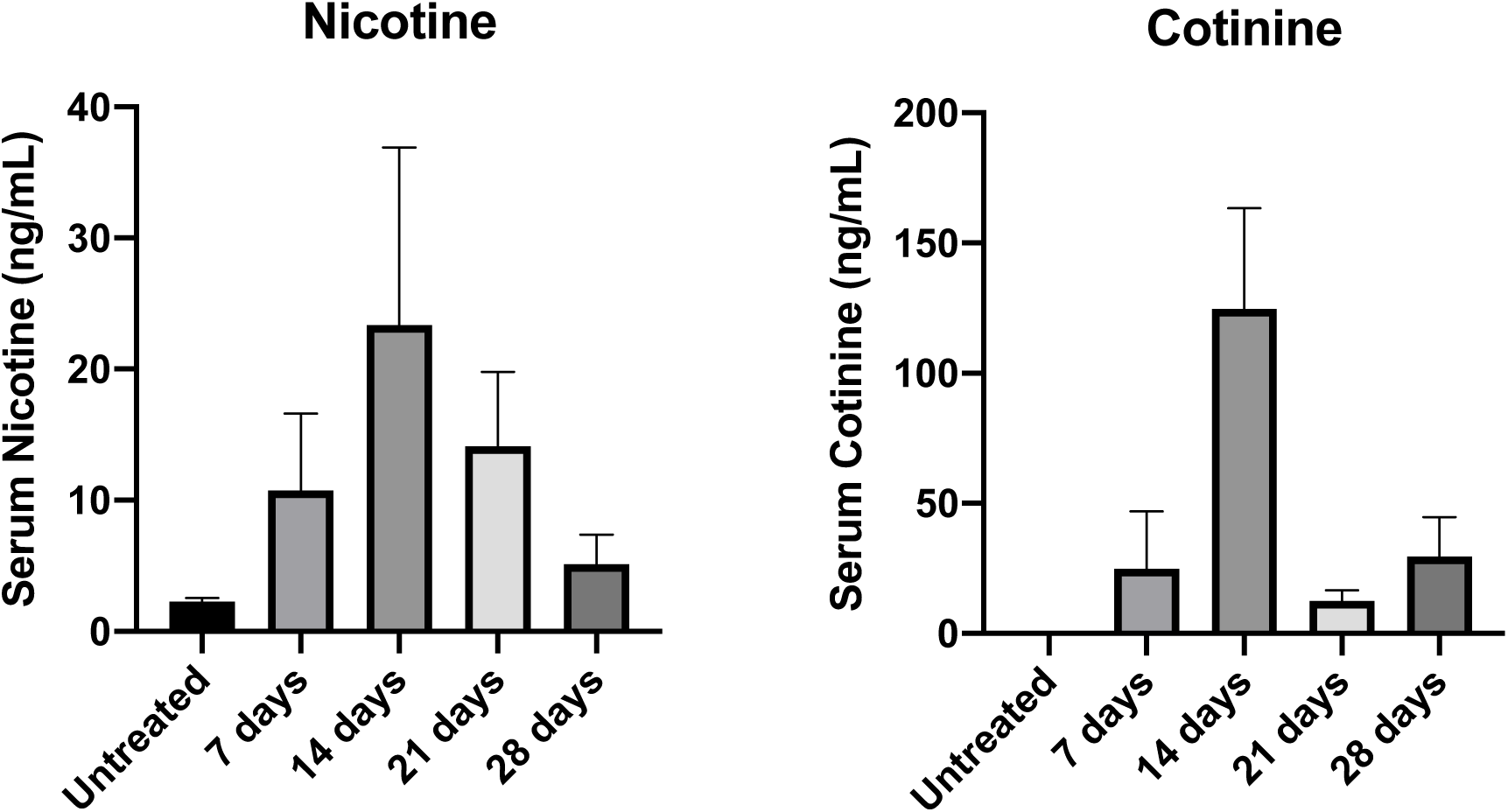
(A)Nicotine and (B) cotinine serum levels in FVB/NJ mice given access to nicotine drinking water (100 μg/ml) or vehicle-treated drinking water (untreated). Average serum levels (ng/ml) from 5 mice collected after 7, 14, 21, or 28 days. Error bars represent standard deviation between mice.

## Supplemental tables

**Supplementary Table 1: Samples included in this study**

Samples analyzed by RNAseq, organized by oncogenic driver and model type. Number of mice and individual tissue samples collected are listed.

**Supplementary Table 2: GEMM v. allograft cluster GSEA**

GSEA analysis of differential gene expression between GEMM and allograft models of metastatic breast cancer.

Sheet 1: Gene ontology terms positively enriched in GEMM v. Allograft GSEA

Sheet 2: Gene ontology terms negatively enriched in GEMM v. Allograft GSEA

Abbreviations: NAME (Gene ontology term), SIZE (Number of genes in the gene set after filtering out those genes not in the expression dataset), ES (enrichment score), NES (normalized enrichment score), NOM p-val (nominal p-value), FDR q-val (false discovery rate), FWER p-val (familywise-error rate), and RANK AT MAX (the position in the ranked list at which the maximum enrichment score occurred).

**Supplementary Table 3: GEMM v. allograft MSGE IPA**

IPA analysis comparing gene expression pathways in GEMM MSGE v. allograft MSGE.

**Supplementary Table 4: Orthotopic v. Tail vein MSGE**

MSGE from orthotopic injection v. tail vein injection models using 4T1 (Sheet 1), 6DT1 (Sheet 2), and Mvt1 (Sheet 3) cell lines.

**Supplementary Table 5: Orthotopic v. Tail vein MSGE IPA**

IPA of the MSGE from orthotopic injection v. tail vein injection models using 4T1 (Sheet 1), 6DT1 (Sheet 2), and Mvt1 (Sheet 3) cell lines.

**Supplementary Table 6: IPA common to spontaneous models of breast cancer metastasis**

IPA of MSGE common to spontaneous models (GEMM and orthotopic injection) of breast cancer metastasis.

**Supplementary Table 7: Sildenafil dosing and human equivalent**

FDA established dose of sildenafil citrate in humans and mice, compared to the dose and serum Cmax levels achieved in this study.

**Supplementary Table 8: Nicotine and cotinine levels as measured by LC-MS/MS**

Nicotine and cotinine serum levels as determined by LC-MS/MS in mice administered nicotine drinking water. Serum collected from five mice once a week for four consecutive weeks.

Sheet 1: Summary table

Sheet 2: Raw data

Abbreviations: %CV (coefficient of variation), Avg (Average), Nic (Nicotine), and Cot (Cotinine).

**Supplementary Table 9: Primer sequences**

List of primers used for qRT-PCR analysis.

